# Bat-borne influenza A H9N2 virus shows species-specific replication and transmission

**DOI:** 10.64898/2026.10.15.759517

**Authors:** Jan Schinköthe, Jacob Schön, Lorenz Ulrich, Andrea Aebischer, Reiner Ulrich, Annika Graaf-Rau, Anne Balkema-Buschmann, Ghazi Kayali, Kevin Ciminski, Martin Schwemmle, Donata Hoffmann, Martin Beer, Nico Joel Halwe

## Abstract

Bat-borne H9N2 IAV (bat H9N2), initially detected in bat feces in Egypt in 2017, has recently attracted attention because of its unexpected pre-pandemic properties, including airborne transmission between ferrets and efficient replication in human lung explant cultures. However, its host range and potential reservoir hosts remain poorly defined, particularly with respect to additional bat hosts and mammalian or avian mixing-vessel species. Here, we compare the susceptibility of two pteropodid bat species, *Rousettus aegyptiacus* and *Eidolon helvum*, and additionally assessed pigs and quails following intranasal inoculation with bat H9N2.

Juvenile *R. aegyptiacus* showed only low-level viral replication and shedding, largely restricted to the upper respiratory tract, and did not show transmission to contact bats. In contrast, both juvenile and adult *E. helvum*, despite showing no clinical signs, showed high-level viral replication and shedding in the upper respiratory tract, with efficient and early transmission to contact bats. Viral RNA was most abundant in the respiratory turbinates, the nasopharyngeal meatus and the tracheal epithelium, where it was associated with moderate to severe necrotizing rhinitis and tracheitis. The lower respiratory tract was less involved, with viral RNA confined to scattered bronchioles and associated with focal necrotizing bronchiolitis. Other organs, including the gastrointestinal tract, lacked histologic lesions and viral RNA labeling. In pigs and quails, intranasal inoculation did not result in detectable replication, transmission, or seroconversion.

Our findings demonstrate marked differences in the susceptibility to bat H9N2 between the two fruit bat species tested and identify *E. helvum* as a highly susceptible host capable of rapid and efficient transmission. Conversely, pigs and quails were not susceptible, suggesting that additional viral adaptations may be required to allow for a productive infection in these species.

**Author Summary:** Bat-borne H9N2 IAV has raised concern because previous studies demonstrated efficient transmission between ferrets, replication in human lung tissue and human MxA-escape under experimental conditions [1]. This suggests that bat H9N2 has several characteristics associated with zoonotic potential. However, it remained unclear which bat species are susceptible to infection and capable of supporting virus transmission. Furthermore, it is unknown whether other mammalian species could serve as reassortment hosts for bat H9N2 with other IAVs.

Here, we experimentally tested the virus in two different Old World bat species, *Eidolon (E.) helvum* and *Rousettus (R.) aegyptiacus*, as well as in pigs and quails: *R. aegyptiacus* showed only low levels of bat H9N2 infection and did not transmit the virus to contact animals. In contrast, *E. helvum* supported high-level replication of bat H9N2 in the upper respiratory tract, also allowing efficient transmission to contact bats. Although subclinical, infection in this species was also associated with marked tissue damage in nose, throat, and trachea. In contrast, experimentally infected pigs and quails showed no evidence of viral replication, transmission, or immune responses. Our findings highlight that susceptibility to bat H9N2 is strongly species-dependent and identify *E. helvum* as a putative bat reservoir host.

## Introduction

Influenza A viruses (IAVs) remain among the most important zoonotic pathogens because their broad host range and capacity for adaptive evolution permit repeated cross-species transmission and, occasionally, pandemic emergence. This concern has intensified during the ongoing H5N1 panzootic and spill-over to mammals, including dairy cattle in the US, underscoring how rapidly IAV ecology can shift once host barriers are breached [2–4]. In parallel, avian-origin H9N2 viruses continue to cause sporadic human infections [5]. To date, up to 195 zoonotic human H9N2 infections have been reported [6], most in children and associated with mild to moderate upper respiratory tract disease [7]. However, severe and even fatal courses have also been described, particularly in individuals with underlying conditions [5,8], highlighting the public health relevance of this subtype.

In 2017, a genetically distinct H9N2 IAV from Egyptian fruit bats *(R. aegyptiacus*) was discovered and successfully isolated [9]. In 2018, bat H9N2 was detected in *R. aegyptiacus* from South Africa [10], suggesting a widespread circulation among African fruit bats. Time-calibrated phylogeny indicated that bat H9N2 emerged as a reassortant, combining internal gene segments of an Old World bat IAV backbone with more recent avian-derived H9 and N2 segments [1].

Previous studies demonstrated that bat H9N2 has a mammalian-like pH stability and neuraminidase activity, and replicates productively in human lung explants, evasion of restriction by human MxA, high virulence in mice and efficient, even airborne, transmission between ferrets [1,11]. MxA escape is a relevant pre-pandemic feature, as this interferon-induced antiviral factor represents a major species barrier for zoonotic IAV [12,13]. Additionally, bat H9N2 receptor binding assays indicate an exclusive α2,3-sialic acid receptor binding, which is rather typical for avian IAV but still enables an efficient transmission between ferrets [1,11].

We previously showed that juvenile to subadult *R. aegyptiacus* bats were barely susceptible to experimental infection with bat H9N2 and that the virus did not transmit to direct contacts [14], suggesting that this species might not represent the only reservoir.

Based on these findings, we addressed three key questions: First, given the broad susceptibility of different species to experimental bat H9N2 infection, we asked whether relevant intermediate hosts are also susceptible. We therefore infected pigs (*Sus scrofa domesticus*) and quails (*Coturnix japonica*), two major influenza A virus mixing-vessels that express both α2,3- and α2,6-linked sialic acid receptors and can support reassortment and cross-species adaptation [15,16]. Second, because inefficient transmission in *R. aegyptiacus* would be expected to limit sustained circulation of bat H9N2, we investigated the potential role of *E. helvum* in the viral ecology, particularly in light of reported H9 seropositivity in this species in Ghana [17]. Third, we examined whether host age influences susceptibility and transmission by including more juvenile *R. aegyptiacus*, similar to age-dependent effects recently described for Marburg-virus (MARV) [18]. Across all species and age groups, we assessed pathogenesis, viral replication, shedding, seroconversion, and transmission to direct contacts.

Our data show that unlike *R. aegyptiacus*, bat H9N2 replicates in and transmits efficiently among *E. helvum*, causing pathological lesions in the upper respiratory tract (URT). On the other hand, we show that pigs and quails are not susceptible to an experimental infection with bat H9N2. Further studies are needed to assess if additional genetic changes can lead to an adaption to these hosts allowing efficient replication and transmission.

## Results

### Limited replication without transmission in juvenile *R. aegyptiacus*

To determine the susceptibility of juvenile *R. aegyptiacus* to infection with bat H9N2 (A/bat/Egypt/381OP/2017), 15 male and female bats between four and twelve months of age were intranasally inoculated with 10^4.96^ TCID_50_ per animal (Supplementary Figure 1A). One day post inoculation, three naïve contact animals were co-housed to monitor transmission (Supplementary Figure 1A). Throughout the course of the experiment none of the inoculated donor *R. aegyptiacus* bats (Supplementary Figure 1B) and none of the contact bats showed any clinical signs or demonstrated any significant body weight loss (Supplementary Fig. 1C). Bat H9N2-specific RT-qPCR analysis of oral swab samples revealed low viral genome copy numbers in the inoculated animals until 7 days post infection (dpi; <10^5^ genome copies mL^-1^) (Fig. 1A), while all contact animals remained negative (Fig. 1B). Consistently, only tissue samples from inoculated animals collected at 6 dpi tested positive for bat H9N2 viral genome mostly in the URT (nasal conchae, larynx/pharynx, trachea) (Fig. 1E. Serological analysis of serum samples showed that inoculated bats mounted a virus-specific antibody response targeting the bat H9 HA, bat N2 NA (Fig. 1F) and the viral NP (Supplementary Fig. 1D) antigens at 21 dpi. In addition, neutralizing antibodies were detected with titers of up to 1:645 (Fig. 1G). Importantly, all contact animals remained sero-negative (Fig. 1F and G).

**Fig 1.**
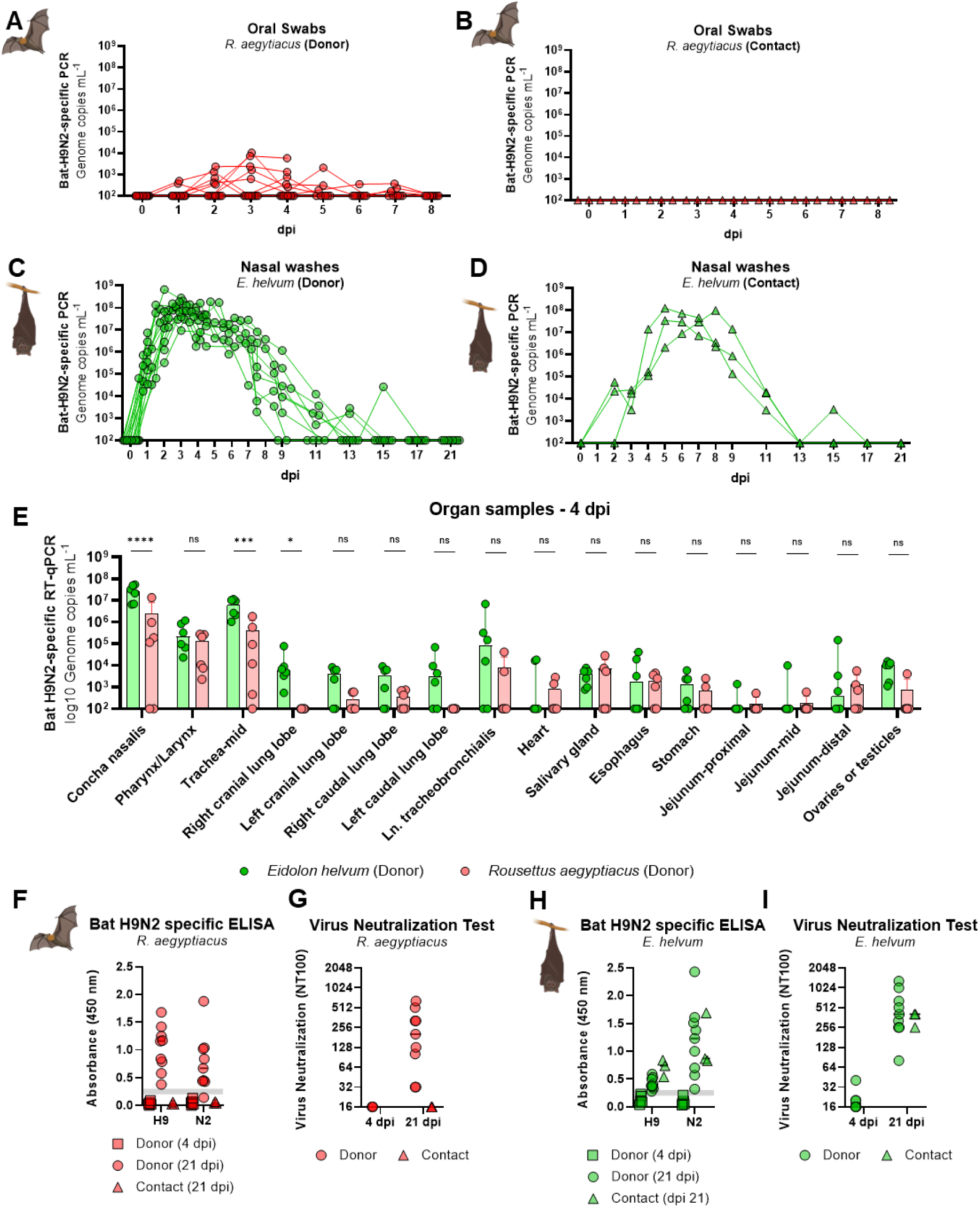
Viral load and humoral immune response after experimental infection with bat H9N2 in *R. aegyptiacus* and *E. helvum.* Experimental infections with bat H9N2 in *R. aegyptiacus* and *E. helvum* were carried out independently. Detailed experimental setup is shown in Supplement Figure 1A. **(A)** Bat H9N2-specific RT-qPCR analysis of oral swabs from inoculated *R. aegyptiacus* (Donor) and **(B)** from non-inoculated contact animals. **(C)** Bat H9N2 specific RT-qPCR analysis of nasal washes of bat H9N2 inoculated *E. helvum* bats (donors) and **(D)** non inoculated contact *E. helvum* bats. **(E)** Bat H9N2-specific RT-qPCR analysis of organ samples from *R. aegyptiacus and E. helvum* donor animals at 4 dpi. Bars indicate median with 95 % CI and statistical comparisons based on the logarithmic values were performed using an ordinary two-way ANOVA followed by Bonferroni’s multiple comparisons, assuming a single pooled variance. ns, not significant; * = *p* < 0.05; *** = *p* < 0.001; **** = *p* < 0.0001. **(F)** Bat-H9 and bat-N2 specific ELISA-results and **(G)** Virus neutralization test (VNT_100_) of donor and contact *R. aegyptiacus* bats. **(H)** Bat-H9 and bat-N2 specific ELISA-results and **(I)** Virus neutralization test (VNT_100_) of donor and contact *E. helvum* bats.

### Efficient viral replication and transmission of bat H9N2 in mixed age *E. helvum*

To determine susceptibility to bat H9N2 infection in *E. helvum*, 15 male and female bats aged four months to three years were inoculated intranasally with 10^4.34^ TCID_50_ per animal (Supplementary Figure 1A). One day post inoculation, three naïve contact bats were co-housed to monitor transmission (Supplementary Figure 1A). Throughout the course of the experiment, none of the inoculated donor bats (Supplementary Figure 1E) and none of the contact bats showed any clinical signs or demonstrated sustained body weight loss (Supplementary Figure 1F). Following experimental inoculation, *E. helvum* bats showed substantial viral shedding between 1 to 15 dpi, with peak viral genome loads in nasal wash samples exceeding 10^8^ genome copies mL^-1^ at 2 - 4 dpi (Fig. 1C). Already 1 day post co-housing (2 dpi), contact bats shed viral genome loads in nasal washing samples comparable to that of the inoculated donor animals (Fig. 1D). Infectious virus titers of up to 10^7^ TCID_50_/mL were detected by endpoint dilution assay in the nasal wash samples of the donors (Supplementary Figure 1G) and also in samples from all contact bats (Supplementary Figure 1H), further confirming efficient virus replication and intraspecies transmission in *E. helvum*.

Necropsy at 4 dpi followed by examination of organ samples revealed that bat H9N2 replication was predominant in the URT, specifically in the nasal conchae, pharynx/larynx and the trachea (Fig. 1E). Interestingly, the bat H9N2 viral genome loads were 10 to 100-fold higher in URT samples from *E. helvum* than in *R. aegyptiacus* (Fig 1E). In the lower respiratory tract (LRT), including the four different lung samples and the tracheobronchial lymph node, much lower viral genome copies than in the URT were detected in both species (Fig. 1E). Nevertheless, *E. helvum* demonstrated a nonsignificant trend towards higher viral loads in tissue samples from the LRT (10^3^ - 10^6^ genome copies mL^-1^) as compared to *R. aegyptiacus* (10^2^ - 10^5^) (Fig. 1E). Moreover, 6/6 donor *E. helvum* bats had detectable bat H9N2 viral genome copies at least in one out of five LRT tissue samples, in contrast to only 3/6 *R. aegyptiacus* donor bats in at least one out of five LRT tissue samples, both at 4 dpi (Fig. 1E). Using RT-qPCR, weakly positive signals for bat H9N2 viral RNA were also observed in gonad/ovary and brain samples of *E. helvum* bats. However, these findings could not be confirmed by ISH. Although productive infection was confirmed by viral genome detection in swab and organ samples, seroconversion was less abundant, but still detectable with antibodies to the viral glycoproteins bat H9 HA and bat N2 NA in ELISA (Fig.1H), and the IAV NP (Supplementary Figure 1I). Neutralizing antibodies were detected by VNT in all donor and contact animals with titers of up to 1:1,290 at 21 dpi (Fig.1I),

These results show that bat H9N2 replication was more efficient in *E. helvum* than in *R. aegyptiacus* fruit bats. Moreover, bat H9N2 is transmissible only between *E. helvum* and not between *R. aegyptiacus* fruit bats in our experimental infection model.

### Replication of bat H9N2 in the URT of *E. helvum* is associated with necrotizing rhinitis

At 4 dpi, marked species differences in URT infection became evident between *E. helvum* and *R. aegyptiacus* (Fig 2A–P). ISH revealed significantly higher bat H9-specific vRNA levels in the respiratory turbinate epithelium of *E. helvum* than *R. aegyptiacus* (Fig 2A). In *E. helvum*, vRNA labeling in the respiratory turbinate epithelium was prominent and often coalescing (Fig 2B, E, F) and corresponded to moderate necrotizing rhinitis (Fig 2C, D), whereas *R. aegyptiacus* showed only absent to minimal labeling in this region with no epithelial necrosis.

**Fig 2.**
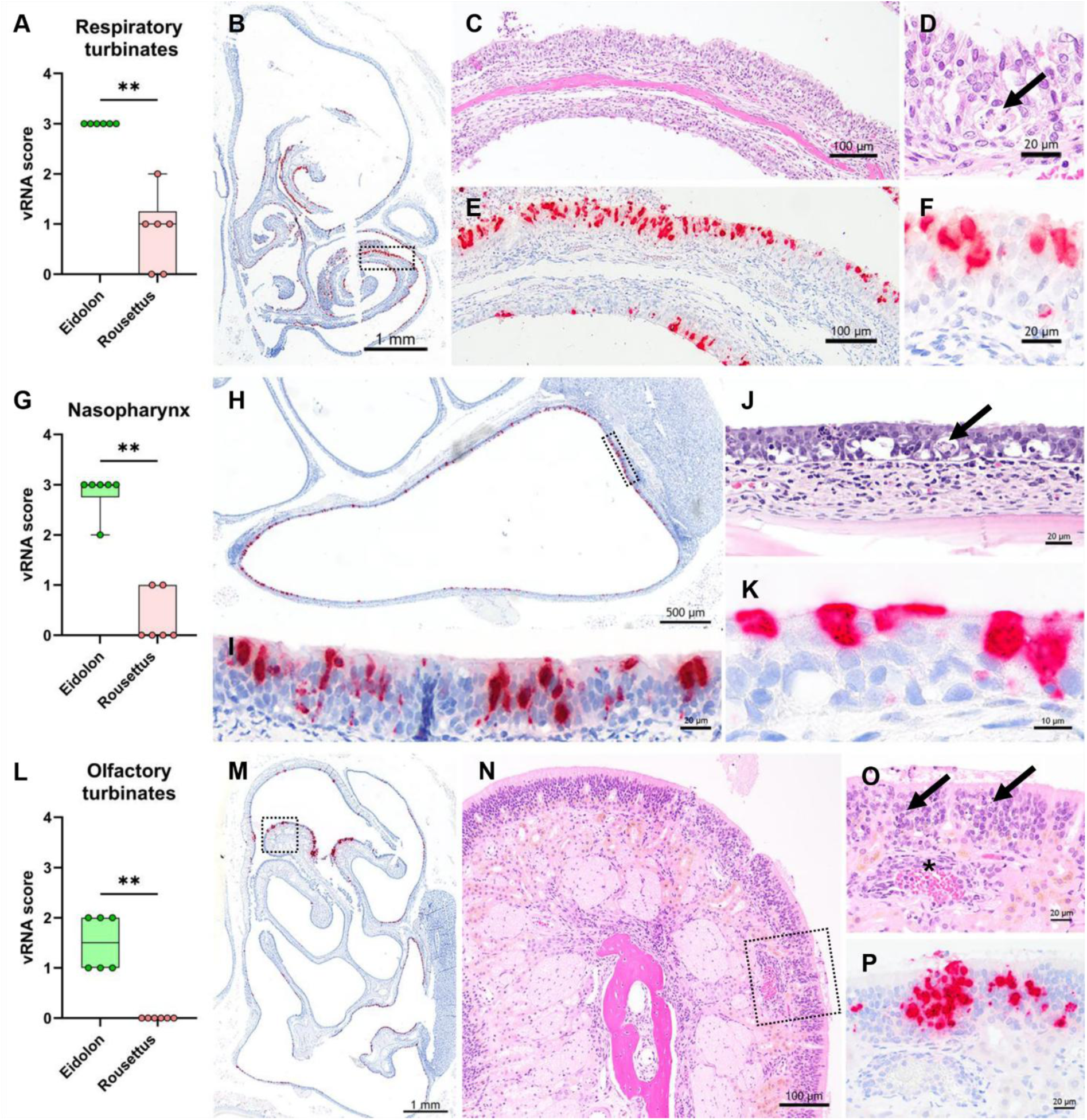
Characterization of lesions by histopathology and bat H9N2-specific vRNA distribution by *in situ* hybridization in distinct anatomical regions of the nasal cavity and nasopharynx of *E. helvum* at 4 dpi. **(A)** Respiratory turbinates at level I show significantly higher vRNA scores in *E. helvum* than in *R. aegyptiacus.* **(B)** Coalescing bat H9N2-specific vRNA labeling is present in the respiratory epithelium lining the nasal turbinates. **(C)** Higher magnification of the boxed area in B shows moderate necrotizing rhinitis characterized by loss of microvilli, epithelial sloughing, and mild neutrophilic inflammation within the lamina propria. **(D)** Higher magnification of the respiratory epithelium highlights necrosis of a small cluster of respiratory epithelial cells (arrow). **(E)** Corresponding consecutive section to C showing coalescing bat H9N2-specific vRNA labeling confined to the respiratory epithelium. **(F)** Higher magnification of bat H9N2-specific vRNA signal in respiratory epithelial cells. **(G)** The respiratory epithelium lining the nasopharyngeal meatus shows significantly higher vRNA scores in *E. helvum* than in *R. aegyptiacus*. **(H)** Focused view of the nasopharyngeal meatus at level III with coalescing bat H9N2-specific vRNA labeling in the respiratory epithelium. **(I)** Higher magnification of the boxed area in H. **(J)** Higher magnification of the respiratory epithelium highlighting multifocal necrosis of small clusters of respiratory epithelial cells (arrow). **(K)** Corresponding consecutive section to J showing detailed bat H9N2-specific vRNA labeling in respiratory epithelial cells. **(L)** Olfactory epithelium shows significantly higher vRNA scores in *E. helvum* than in *R. aegyptiacus*. **(M)** At level II, the few olfactory turbinates present are multifocally positive for bat H9-specific vRNA. **(N)** Overview of the multilayered olfactory mucosa with underlying lightly brown-pigmented Bowman’s glands and fila olfactoria. **(O)** Higher magnification of the boxed area in N showing multifocal single-cell necrosis (arrows) and prominent endothelial hypertrophy in a medium-sized vessel (asterisk), accompanied by hyperemia and mild perivascular diapedesis of neutrophils and mononuclear cells. **(P)** Corresponding consecutive section to O showing patchy, multifocal bat H9N2-specific vRNA labeling confined to the olfactory epithelium in areas with single-cell necrosis. **A, G, L** Box-and-whisker plots show the median, interquartile range, and minimum-to-maximum range; individual symbols represent single animals (n=6 per group). Statistical comparisons were performed using a two-tailed Mann–Whitney U test. ** = p < 0.01. C, D, J, N, O hematoxylin & eosin staining. B, E, H, I, K, M, P *in situ* hybridization, bat H9N2-specific vRNA, Fast Red chromogen (red), hematoxylin counterstain (blue).

A similar pattern was observed in the nasopharynx. The respiratory epithelium lining the nasopharyngeal meatus of *E. helvum* had significantly higher vRNA scores than in *R. aegyptiacus* (Fig 2G). In *E. helvum*, viral RNA was distributed in a coalescing pattern along the epithelium (Fig 2H, I, K) and was associated with multifocal necrosis of small groups of respiratory epithelial cells (Fig 2J). In contrast, *R. aegyptiacus* did not develop comparable lesions, and vRNA scores in this region remained restricted to 0 or 1.

The olfactory region showed the same species divergence. Olfactory turbinates of *E. helvum* had significantly higher vRNA scores than those of *R. aegyptiacus* (Fig 2L). ISH-signals in the olfactory epithelium of *E. helvum* were mostly multifocal and clustered (Fig 2M), and affected areas exhibited multifocal single-cell necrosis together with vascular signs of acute inflammation, including endothelial cell swelling, hyperemia, and mild perivascular leukocyte diapedesis (Fig 2N, O). The corresponding consecutive section demonstrated patchy, multifocal bat H9-specific vRNA labeling confined to the olfactory epithelium (Fig 2P). vRNA labeling was absent or focal and only rarely single cell epithelial necrosis was evident in the olfactory epithelium at levels III and IV of the nasal cavity. In contrast, no vRNA was detected in olfactory turbinates of *R. aegyptiacus,* and consequently, no epithelial necrosis was observed.

In the Waldeyer’s ring, the palatine tonsil showed significantly increased vRNA scores, mostly as scant intracytoplasmic signals in round to spindloid mononuclear cells (interpreted as dendritic cells or macrophages) in *E. helvum*, when compared to *R. aegyptiacus* (Supplemental Figure 3B) without signs of epithelial degeneration or necrosis of the follicle associated epithelium.

Collectively, these findings indicate that productive epithelial infection with associated moderate necrotizing lesions was a feature of *E. helvum*, whereas infection in *R. aegyptiacus* remained sparse and was not associated with histologically evident necrotizing rhinitis.

### Replication of bat H9N2 in the LRT of *E. helvum* is confined to the tracheobronchial epithelium

At 4 dpi, the LRT of *E. helvum* showed prominent bat H9N2 infection in the trachea but only limited involvement of the lung parenchyma (Fig 3A–G). In the trachea, *E. helvum* had significantly higher vRNA scores than *R. aegyptiacus* (Fig 3A). In infected *E. helvum*, the tracheal mucosa exhibited moderate to severe necrotizing tracheitis with epithelial denudation extending to the basement membrane (Fig 3B), and diffuse bat H9N2-specific vRNA labeling of the tracheal epithelium was present in corresponding sections (Fig 3C). Widespread sloughing of necrotic, vRNA-positive epithelial cells into the lumen was evident, whereas luminal blood was interpreted as a necropsy-related artifact (Fig 3B, C).

**Fig 3.**
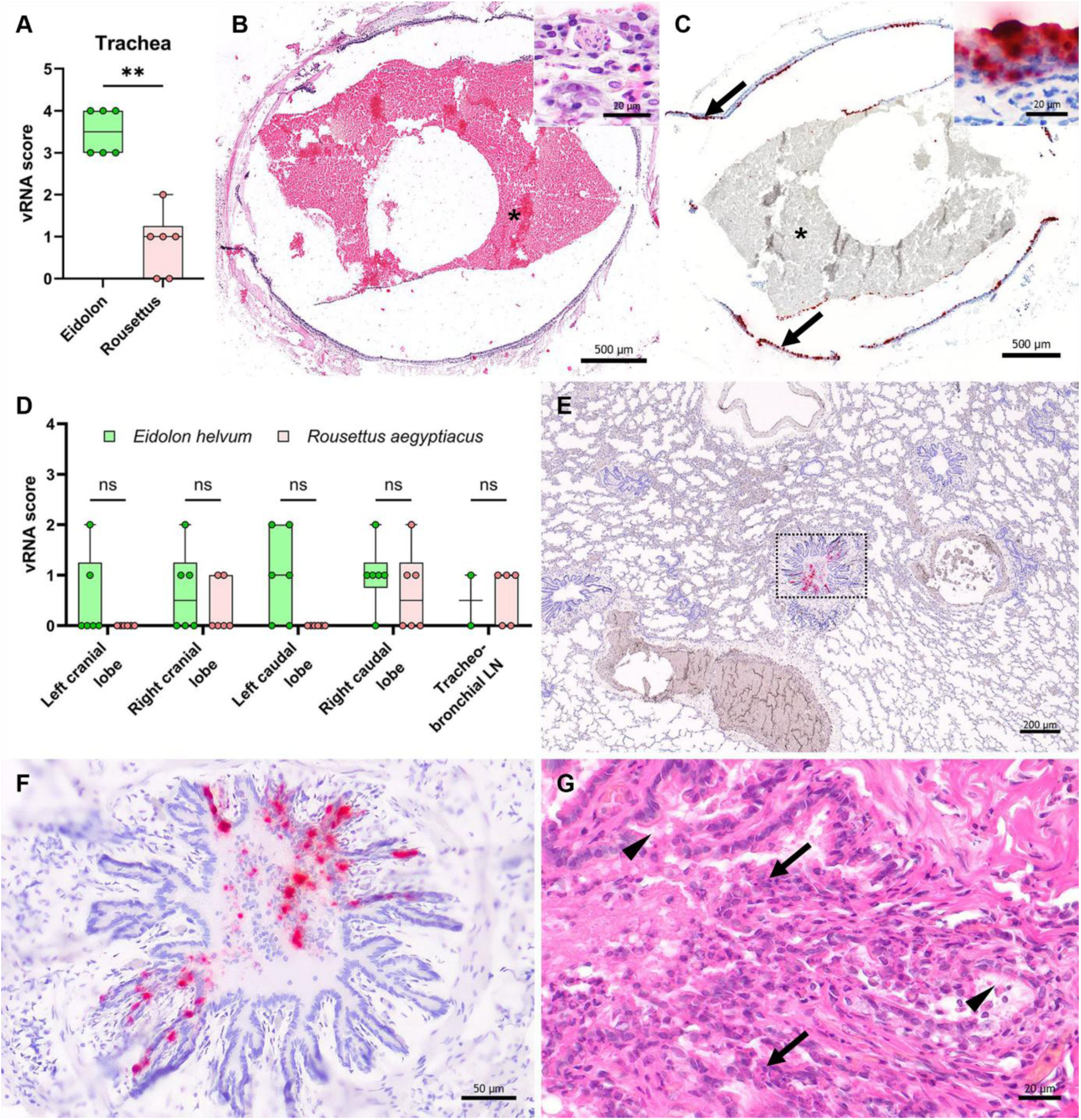
Characterization of lesions by histopathology and bat H9N2-specific vRNA distribution by *in situ* hybridization in distinct anatomical regions of the LRT of *E. helvum* at 4 dpi. **(A)** The trachea shows significantly higher vRNA scores in *E. helvum* than in *R. aegyptiacus*. Box-and-whisker plots show the median, interquartile range, and minimum-to-maximum range; individual symbols represent single animals (n=6). Statistical comparisons were performed using a two-tailed Mann–Whitney U test. ** = p < 0.01. **(B)** Moderate to severe necrotizing tracheitis with epithelial denudation extending to the basement membrane. The asterisk indicates extravasated blood within the tracheal lumen, interpreted as a necropsy-associated artifact. **(C)** The tracheal respiratory epithelium is diffusely positive for bat H9N2-specific vRNA, with widespread sloughing of necrotic, vRNA-positive epithelial cells into the lumen and onto the extravasated blood (asterisk). The inset shows diffuse vRNA labeling of the tracheal epithelium at higher magnification. **(D)** In the lung, bat H9N2-specific vRNA was restricted to single or few bronchioles within the examined lung lobes. No significant differences were observed between bat species or among lung lobes. Bat H9N2-specific vRNA was also detected in some draining tracheobronchial lymph nodes (Tracheo-LN). Box-and-whisker plots show the median, interquartile range, and minimum-to-maximum range; individual symbols represent single animals (n = 2–6 per group). Species were compared separately at each anatomical site using two-tailed Mann–Whitney U tests. ns, not significant. **(E)** Left caudal lung lobe shown as an example of the pulmonary distribution pattern, with vRNA signals distributed randomly and largely confined to single bronchioles. **(F)** Higher magnification of the boxed area in **E**, showing multifocal bat H9N2-specific vRNA labeling in bronchiolar respiratory epithelium and luminal debris. **(G)** Consecutive section of **E** showing moderate necrotizing bronchiolitis with abundant necrotic luminal debris (arrows) and only a few remaining intact respiratory epithelial cells with preserved kinocilia (arrowheads). **(B, G)** Hematoxylin and eosin stain. **(C–F)** In situ hybridization for bat H9-specific vRNA with Fast Red chromogen (red) and hematoxylin counterstain (blue).

In contrast to the trachea, pulmonary involvement was very limited. Across the examined lung lobes, bat H9N2-specific vRNA was detected only in single or few bronchioles and showed no significant differences between species or among lung lobes (Fig 3D). Similar sparse labeling was also observed in some round to spindloid mononuclear cells (interpreted as dendritic cells or macrophages) in the draining tracheobronchial lymph nodes. In the lung, vRNA signals were randomly distributed and largely confined to bronchiolar epithelium and luminal debris (Fig 3E, F). Corresponding histologic sections showed moderate necrotizing bronchiolitis with abundant necrotic intraluminal debris and only few remaining intact epithelial cells retaining kinocilia (Fig 3G).

Collectively, these findings indicate that in *E. helvum*, bat H9N2 infection of the LRT was dominated by severe tracheal epithelial involvement, whereas pulmonary lesions were focal and manifested as necrotizing bronchiolitis. No histologic lesions or bat H9N2-specific vRNA labeling were detected in the gastrointestinal tract or in spleen, liver, kidney, and brain.

### Quails (*Coturnix japonica*) are not susceptible to experimental infection with bat H9N2

To assess the susceptibility of quails (*Coturnix japonica*), a possible IAV mixing vessel, we inoculated 15 quails of mixed sexes and more than three weeks of age with 10^4.94^ TCID_50_ per animal of bat H9N2 (Supplement Figure 4A). To monitor transmission, naïve contact quails were co-housed at 1 dpi. Throughout the course of the experiment, no clinical signs or body weight loss were observed in inoculated quails (Fig. 4A) and naïve contact quails (Fig. 4B). RT-qPCR analysis of oropharyngeal swabs from experimentally infected quails revealed only trace amounts of viral genome in 8/15 inoculated animals at variable timepoints up to 6 dpi, with peak viral load of <10^5^ genome copies ml^-1^ (Fig. 4C). None of the swab samples from the contact animals tested positive for bat H9N2 viral genome at any of the evaluated timepoints (1 – 6 dpi) (Fig. 4D). Evaluation of pharynx/larynx samples for bat H9N2 genome revealed viral genome traces only for 1/6 samples (Fig. 4E). All the other 13 evaluated organ/tissue samples (nasal concha, mid-trachea, left lung, right lung, heart, proventriculus, gizzard, jejunum, cecum, colon, liver, kidney, spleen) tested negative for viral genome (Supplementary Figure 4B). Consistently, none of the inoculated or naïve contact quails seroconvert at 21 dpi (Fig. 4F). Together these results indicate that quails show minimal to no susceptibility to intranasal inoculation with bat H9N2 under experimental conditions.

**Figure 4.**
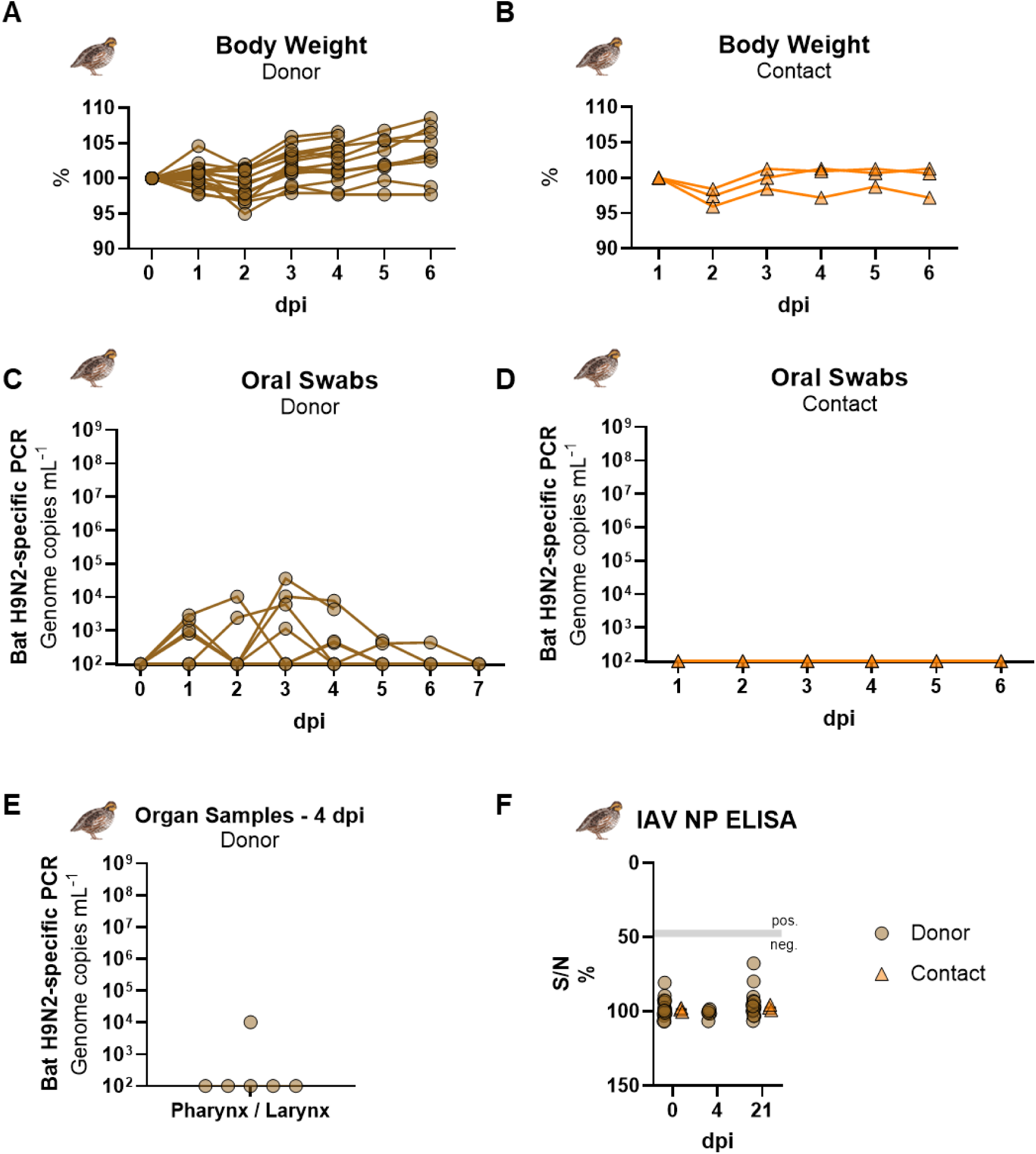
Inoculation of quails (*Coturnix japonica*) with bat H9N2. **(A)** Body weight of bat H9N2-infected donor quails and **(B)** direct contact quails over time. **(C)** Bat H9N2-specific RT-qPCR analysis of oropharyngeal swabs from infected donor quails and **(D)** direct contact quails. **(E)** Only one animal had minimal amounts of viral RNA in the pharynx/larynx sample. All other organs in all other six animals euthanized at 4 dpi were negative for bat H9N2 viral RNA (Supplement Figure 4B). **(F)** Serum samples were evaluated by IAV NP-specific ELISA.

### Pigs (*Sus scrofa domesticus*) do not support bat H9N2 replication and transmission

Given the role of pigs as the principal mixing vessel for avian and mammalian IAVs, we assessed their susceptibility to experimental infection with bat H9N2. We intranasally inoculated nine pigs with 10^4.32^ TCID_50_ per animal and monitored virus transmission to naïve contact pigs from 1 dpi onwards (Supplementary Figure 4A). Throughout the course of the experiment, no clinical signs or elevated body temperature were observed in inoculated donors (Fig. 5A) and contact pigs (Fig. 5B). RT-qPCR analysis of the nasal swab samples revealed only minimal detection of bat H9N2 viral genome in 3/9 directly inoculated animals (peak viral load <10^4^ genome copies ml^-1^ (Fig 5C)). In line with that, none of the nasal swabs of the contact animals tested positive for bat H9N2 viral genome at any of the evaluated time points (0 – 8 dpi) (Fig. 5D). Only 1/6 post-mortem bronchial swabs from 8 dpi necropsies tested positive for bat H9N2 viral genome, with <10^4^ genome copies ml^-1^ (Fig. 5E). At total of 22 organ/tissue samples of different sites (URT, LRT, gastrointestinal tract, gut-associated lymphoid tissue, spleen, parenchymal organs and brain) were taken at necropsy and all tested negative for bat H9N2 viral genome (Supplementary Figure 4C). All donor and contact pigs remained seronegative at all specific necropsy timepoints (2,4,8 or 21 dpi) as tested by the IAV-NP specific ELISA (Fig. 5F). Collectively, the pig model demonstrated no susceptibility to experimental infection with bat H9N2.

**Figure 5.**
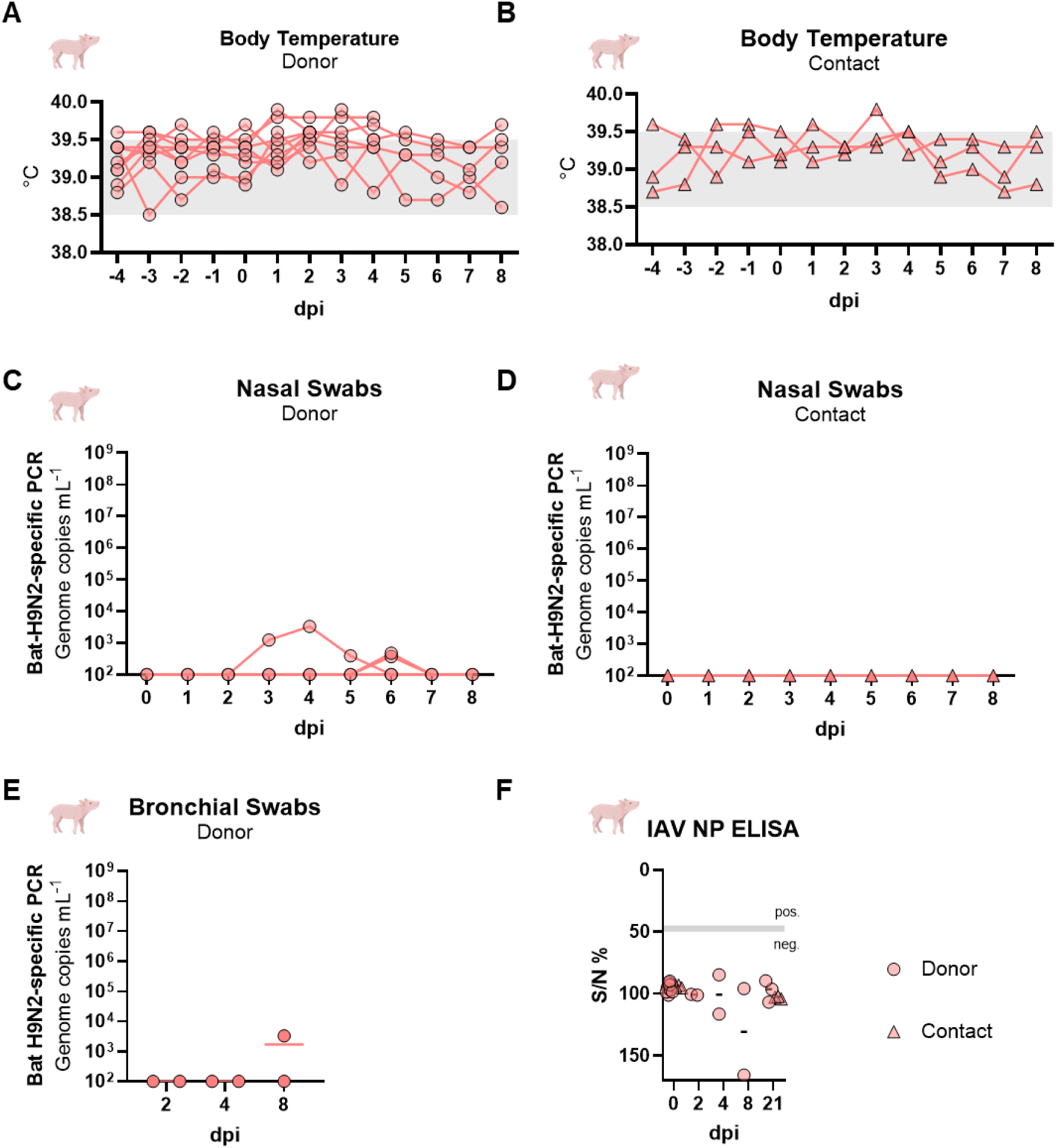
Experimental infection of pigs (*Sus scrofa dom.*) with bat H9N2. **(A)** Body temperature of bat H9N2-inoculated donor pigs and **(B)** direct contact animals over time. Thermoneutral condition range is indicated as grey area. **(C)** Bat H9N2-specific RT-qPCR analysis of nasal swabs from donor pigs and **(D)** contact animals. **(E)** Bat H9N2-specific RT-qPCR analysis of bronchial swabs from infected pigs at 2-, 4- or 8-dpi. Organ samples of pigs were entirely negative for bat H9N2 viral genome (Supplement Figure 4C). **(F)** Serum samples were evaluated by IAV NP-specific ELISA.

## Discussion

Since the discovery of bat H9N2 IAV in 2017 [9], studies of bat H9N2 have broadened our understanding of IAV host range and transmission. Recently, our group and others have uncovered several unexpected pre-pandemic characteristics of bat H9N2, including the ability to infect human lung explants efficiently, evade human MxA restriction, and transmit efficiently between ferrets [1,11]. Furthermore, despite an exclusive receptor binding affinity to α2,3 sialic acid receptors, bat H9N2 also demonstrates a high replication efficiency in the ferret model [1,11]. These results suggest that bat H9N2 is well adapted to the sialic acid repertoire of ferrets. The ability to replicate in mammalian species offers multiple possibilities for reassortment events between bat H9N2 and other avian or human IAVs [3,19–21]. Therefore, characterizing the experimental susceptibility of potential bat H9N2 host species is urgently needed. Only two independent genomic sequences of bat H9N2 are available to date, both originating from *R. aegyptiacus* (Egypt and South Africa) [9,10]. Based on these findings, *R. aegyptiacus* was thought to be the reservoir host of bat H9N2. In the same setup, we demonstrate for the first time that only *E. helvum* fruit bats efficiently replicate and transmit bat H9N2 after intranasal inoculation. In contrast, despite being susceptible to bat H9N2 infection, viral replication was limited and transmission could not be observed in juvenile *R. aegyptiacus*. These results are consistent with our first bat H9N2 experimental infection trial with *R. aegyptiacus* [14], where transmission was also not observed. The lack of bat H9N2 transmissibility in *R. aegyptiacus* bats strongly supports the hypothesis that our observations are not dependent on age, although this has been shown to play an important role within e.g. Marburg virus (MARV)-transmission dynamics [18]. In contrast, our data identify *E. helvum* as a markedly more susceptible host for bat-borne H9N2 IAV than *R. aegyptiacus* and show that this difference is biologically consequential at multiple levels: Viral replication, tissue tropism, lesion severity, intraspecies transmission and seroconversion of contact animals. In *E. helvum*, infection was most prominent in the URT with high viral burdens. In addition, the respiratory turbinate epithelium, the nasopharyngeal meatus, and the tracheal epithelium displayed moderate to severe necrotizing rhinitis, pharyngitis and tracheitis. In contrast, the LRT involvement was limited to scattered bronchioles exhibiting bronchiolitis. By contrast, infection in *R. aegyptiacus* remained sparse, with negative or focal/oligofocal vRNA distribution and without associated lesions or transmission to contact animals. Together, the findings of our experimental infection study suggest that, in addition to *R. aegyptiacus*, *E. helvum* might be a putative reservoir host for bat H9N2. Our experimental data indicates that *E. helvum* may represent the main reservoir host of bat H9N2 allowing transmission, assigning *E. helvum* a potential epidemiological relevance towards bat H9N2 circulation. On the other hand, *R. aegyptiacus* may be a more rare and less efficient spillover host with moderate susceptibility.

The repeated detection of bat H9N2 viral RNA in *R. aegyptiacus*, but not in other bat species, in several regions across Africa may be due to sampling bias [22], particularly following the initial detection of bat H9N2 in this species [9,10]. Thus, serosurveys of *E.helvum* specifically targeting bat H9N2 should be initiated to investigate the seroprevalence of bat H9N2 in this species in the field. In addition, possible scenarios of continued bat H9N2 circulation should involve newborn *R. aegyptiacus* bats with waning maternal immunity, or strong seasonal dependence of viral circulation, as seen for MARV [23] or coronaviruses [24]. Future investigations should therefore focus on how bat H9N2 maintains circulation within *R. aegyptiacus* populations in Africa. Additionally, *E. helvum* and *R. aegyptiacus* are genetically not closely related [25] and favor different roosting habitats – trees and caves, respectively. Despite their distinct distribution patterns, the ranges of both species overlap across substantial parts of sub-Saharan Africa [26,27]. While *R. aegyptiacus* bats are primarily a resident species and just travel up to a maximum of <10 km per night on average, *E. helvum* is known to migrate several hundred kilometers – up to over 2.000 km in single cases – to follow fruit abundance [28]. Therefore, *E. helvum* and *R. aegyptiacus* may occasionally use the same foraging resources. Feeding on the same fruit trees creates an interface for cross-bat viral spillover and may warrant continued bat H9N2 circulation in fruit bats across the African continent.

Fruits as potential mediators of transmission might not only be an interface of bat-bat transmission but may even provide a spillover platform to other mammals, species or even humans. In line with that, African households have been shown to feed their pigs, poultry and other livestock animals with fruits as food and vitamin source [29,30]. Pig and quail farming is popular in certain African countries, especially in central or Southern Africa [31]. Pigs and quails are often discussed as reassortment vessels between all kinds of IAVs, since their respiratory tract harbors both α2,3 and α2,6 sialic acid receptors [32–36]. Importantly, we were able to show here that neither pigs nor quails are susceptible to bat H9N2 after experimental intranasal inoculation. Therefore, it appears that bat H9N2 has to overcome further genetic barriers to establish a productive infection in pigs and quails. The molecular restricting determinants are yet to be determined and will be part of future investigations.

A second major implication of our work is that bat H9N2 behaves very differently from H18N11 at the tissue level. H18N11 has emerged as a useful model for understanding unconventional bat IAV biology, including MHC-II-dependent entry, intestinal replication, and leukocyte tropism, particularly macrophages, B cells, and NK/T cells [37–40]. In our analysis, bat H9-specific vRNA labeling was confined to the mucosal surface, with no evidence of productive infection in round-cell populations within the lamina propria or deeper tissues. The sparse vRNA-positive cells observed in the palatine tonsil and draining respiratory lymph nodes were consistently located in or adjacent to follicles and interfollicular zones and were morphologically most compatible with antigen-processing cells such as dendritic cells or macrophages. We therefore interpret these signals as more likely reflecting uptake and processing of viral material within local adaptive immune compartments rather than true infection of lymphoid cells. In *A. jamaicensis*, H18N11 infection was predominantly associated with the oral mucosa and gut-associated lymphoid tissues, especially Waldeyer’s ring and Peyer’s patches, and was mainly localized to the associated round-cell populations within the inductive and effector compartments of the mucosal immune system [38]. Therefore, in the current study, we comprehensively screened the gastrointestinal tract in both *E. helvum* and *R. aegyptiacus* bat species. While RT-qPCR of gastrointestinal tissues detected low to moderate viral RNA levels, no vRNA was localized within gastrointestinal tissues by *in situ* hybridization. This strongly suggests that the detected viral RNA originated from swallowed URT secretions and was passively carried within the ingesta, rather than representing productive infection of M cells or enterocytes.

By contrast, productive infection in *E. helvum* was centered on the respiratory epithelium, with the greatest viral burden in the URT, where infection was associated with moderate necrotizing lesions. The most pronounced epithelial necrosis occurred in the middle trachea, which also exhibited the highest vRNA scores in *E. helvum*. Although this tropism extended into the LRT, it remained limited to scattered individual bronchioles. This observation is particularly noteworthy because it indicates a distinct degree of LRT susceptibility, which has previously been described in experimental H18N11 infection of *C. perspicillata* [39], but not in *A. jamaicensis* or in bat H9N2-infected *R. aegyptiacus* [14]. This distinction matters because epithelial tropism and shedding of virus laden epithelial debris in the respiratory tract are much more reminiscent of conventional IAV pathogenesis and most likely facilitates airborne or mucosal droplet driven transmission. In summary, the distribution of lesions and vRNA in *E. helvum* is in general agreement with a pathogenetic model comparable to seasonal IAV in other mammals [41].

At present, our study cannot resolve the functional basis for the marked difference in susceptibility between *E. helvum* and *R. aegyptiacus*, including potential contributions of receptor distribution, epithelial host factors, local innate immune responses, or differences in viral clearance. These unresolved questions define an important next step for bat IAV research.

## Conclusion

In summary, bat H9N2 did not establish productive infection in pigs or quails under the conditions tested in this study. However, IAVs are well known for their capacity to adapt rapidly to new hosts and environments, underscoring the need for continued surveillance of bat H9N2. In this context, our findings identify *E. helvum* as a putative reservoir and indicate that future surveillance efforts should extend beyond *R. aegyptiacus* to include *E. helvum* as a priority target. Expanded sequence data from bat H9N2 detected in free-ranging bat populations will be essential to monitor viral evolution, assess genetic stability, and better define the zoonotic potential of this emerging bat IAV.

## Methods

### Virus

The bat-derived H9N2 A/bat/Egypt/381OP/2017 isolate (GenBank accession numbers: MH376902 to MH376909), was propagated in embryonated SPF-chicken eggs for 5 days at 37 °C. Subsequently, the allantoic fluid was harvested, propagated once more on MDCKII-cells (Collection of Cell Lines in Veterinary Medicine CCLV RIE1061) and was subsequently used as virus stock.

### Cells

Madin-Darby Canine Kidney (MDCK) type II cells (CCLV RIE1061) were used. Cells were incubated at 37 °C under 5% CO_2_ atmosphere using a mixture of Minimum Essential Medium (MEM) formulated as a mixture of MEM with Hank’s salts (5,32 g L^-1^) and MEM with Earle’s salts (4,76 g L^-1^), 2 mM L-Gln, nonessential amino acids, adjusted to 850 mg L^−1^ NaHCO_3_, 120 mg L^-1^ sodium pyruvate, pH 7.2 with 10% fetal calf serum (FCS; Bio & Sell GmbH) or without FCS in the presence of tosyl phenylalanyl chloromethyl ketone (TPCK)-treated trypsin (Sigma) after virus addition. Expi293 cells (Thermo Fisher Scientific, Waltham, Massachusetts, U.S.) were grown in Expi293 expression medium (Thermo Fisher Scientific) and polycarbonate Erlenmeyer flasks (Corning, New York, U.S.) at 37°C, 8% CO2, 125 rpm.

### Recombinant protein cloning and expression

The full-length bat-H9 (amino acids 18 - 515 (MH376902.1)) and bat-N2 (amino acids 30 - 461 (MH376904.1)) were amplified from codon-optimized synthetic genes (GeneArt, Regensburg, Germany). The constructs were cloned in the expression vector pEXPR103 (iba lifesciences, Goettingen, Germany) in frame with an N-terminal modified mouse Ig kappa light chain signal peptide and a c-terminal FLAG-tag.

The recombinant proteins were expressed in Expi293 cells. Transfection was performed using the ExpiFectamine293 transfection kit (Thermo Fisher Scientific) according to the manufactureŕs instructions and the cell culture supernatant was harvested five days after transfection. The proteins were purified from the supernatant using ANTI-FLAG® M2 Affinity Gel (A2220; SigmaAldrich/Merck, St. Louis, Missouri, U.S.) as recommended by the manufacturer and eluted with 100 µg/ml FLAG® peptide (Cat. No. F3290; SigmaAldrich/Merck, St. Louis, Missouri, U.S.). Subsequently, the peptide was removed from the preparation by dialysis against an excess of 1X PBS, pH 7.4.

### Animal experiments ethics declarations

All animal experiments with *E. helvum*, *R. aegyptiacus*, pigs and quails were evaluated by the responsible ethics committee of the State Office of Agriculture, Food Safety, and Fishery in Mecklenburg–Western Pomerania (LALLF M-V) and gained governmental approval under the registration numbers LVL MV TSD/7221.3-1-027/23 and 7221.3-1-007/24.

### Experimental challenge of pigs

Twelve pigs of mixed gender in an age range of 10-15 weeks were used in the experiment. Three groups of four pigs (three donor pigs and one contact pig per stable) were kept in three individual and strictly separated, large animal stables under BSL-3 conditions. All pigs were investigated daily for clinical signs. Before inoculation, blood and swab samples of all pigs were taken and serum tested negative by ELISA for bat H9N2-specific antibodies and tested negative via RT-qPCR for bat H9N2 viral genome. Moreover, the three contact pigs were transferred into a further strictly separated animal stable until 24 hours after inoculation of the donor pigs. Subsequently, all 9 donor pigs were inoculated intranasally using a nasal nebulizer device with 10^4.32^ TCID_50_ per animal of bat H9N2 IAV in a 2 mL volume. 1 dpi, the respective contact pigs were co-housed to the donor stables again (donor 3:1 contact setup). All pigs were sampled via nasal swabs for eight consecutive days and at the endpoint of the experiment at 21 dpi. At timepoints 2, 4 and 8 dpi, two donor pigs were euthanized to study the viral organ load in these animals during a previously estimated and probable acute phase of infection. The residual animals were kept until 21 dpi to allow for seroconversion.

### Experimental challenge of quails

Eighteen quails of mixed gender and older than 21 days were used in the experiment. All animals were kept in an aviary (208,8 x 386 x 203 cm (width x depth x height)) under BSL-3 conditions. All quails were sampled (oral swab and blood) and afterwards tested negative for bat H9N2-related antibodies via ELISA and viral genome via bat H9N2-specific RT-qPCR two days prior to infection. Three contact quails were strictly separated from the donor animals immediately before inoculation of the donor quails for 24 hours. Afterwards, 15 donor quails were intranasally inoculated with 10^4.94^ TCID_50_ bat H9N2 IAV per animal (total volume 0.1 mL), distributed evenly into each nostril. At 1 dpi, the uninfected contact quails were co-housed to the donor animals again. Overall condition of the animals was monitored daily, and body weight was measured regularly. Furthermore, all quails were sampled daily via oral swabs for eight days and one last time at 21 dpi. Six donor quails were euthanized at 4 dpi to study the viral organ tropism. The residual animals were kept until 21 dpi to allow for seroconversion.

### Experimental infection of *E. helvum* and *R. aegyptiacus*

Eighteen *E. helvum* (>6 months up to 104 months) and 18 *R. aegyptiacus* (>4 months up to 12 months) of mixed gender were used in this study in two independent experiments (Supplementary Table 1). Age classes for *E. helvum* are defined as follows: juvenile (2 - 6 months), subadult (6 - 24 months) or adult (>24 months) [42]. Age classes for *R. aegyptiacus* are defined as follows: Juvenile (<12 months), subadult (12 - 24 months), adult (>24 months) [43]. Brief isoflurane induced anesthesia was used before inoculation and blood sampling. All animals were tested negative for bat H9N2-related antibodies via ELISA and viral genome via bat H9N2-specific RT-qPCR 2 days prior to infection. Subsequently, 15 *E. helvum* were inoculated intranasally with 10^4.34^ TCID_50_ of bat H9N2 IAV per animal, and 15 *R. aegyptiacus* were inoculated with 10^4.9^ TCID_50_ of bat H9N2 IAV per animal, distributed evenly into each nostril (total volume 0.1 mL). Three animals of both species were strictly separated for 24 hours before inoculation and served as contact animals. The contact animals were co-housed again afterwards at 1 dpi to all inoculated donor animals (15:3 setup per experiment). All animals were evaluated daily for clinical signs, and body weight was measured regularly throughout the experiments. Longitudinal sampling was performed on all animals until no bat H9N2 viral genome was detected via RT-qPCR. *R. aegyptiacus* were sampled via oral swabs, and *E. helvum* were sampled via nasal washings. Nasal washings were only performed on *E. helvum* due to their larger body size. Nasal wash samples were collected under brief isoflurane anesthesia by flushing each nostril with 250 µl phosphate-buffered saline (PBS). Unfortunately, the small body size including the nose of the *R. aegyptiacus* did not allow nasal washings in these animals. At 4 dpi, six donor animals from each species in each experiment were scheduled for euthanasia to study the viral tropism in organ samples in the early phase of infection. The animal trial was terminated at 21 dpi to allow for seroconversion of all residual animals from both species. Of all euthanized bats various organs were sampled immediately after death for subsequent virus detection via RT-qPCR. The remaining bat carcass was fully submerged in appropriate volumes of neutrally buffered 4 % formaldehyde to allow fixation of tissues for up to 3 weeks.

### Sample work-up and analysis

Organ samples (∼1cm^3^ in size) were first homogenized in a 2mL Eppendorf-tube containing 1 mL of Hank’s balanced salts MEM and Earle’s balanced salts MEM (2 mM L-glutamine, 850 mg L^−1^ NaHCO_3_, 120 mg L^−1^ sodium pyruvate, and 1% penicillin–streptomycin) at 300 Hz using a TissueLyser II (Qiagen, Hilden, Germany). The swabs were transferred to 2 mL of cell culture medium containing 1% Baytril (Bayer, Leverkusen, Germany), 0.5% Lincomycin (WDT, Garbsen, Germany) and 0.2% Amphotericin/Gentamycin (Fisher Scientific Waltham, MA, USA), followed by 30 min. incubation at room temperature under roughly shaking at 300 rpm (KS 260 control, IKA, Staufen, Germany). Nasal washes were performed by administering 250 µL of Phosphate buffered saline (PBS) into each nostril of *E. helvum* and collecting the efflux in a 2 mL Eppendorf-tube.

From all samples, 100 µL was extracted via the NucleoMag Vet kit (Macherey&Nagel, Düren, Germany) according to the manufacturer’s instructions on a Biosprint 96 platform (Qiagen). Viral RNA was detected by using bat H9N2-specific primer and probes [14]. Absolute quantification was done using a standard of known concentrations, corresponding to the RNA of the original virus used for inoculation. Quantification was established by the QX200 Droplet Digital PCR System in combination with the 1-Step RT-ddPCR Advanced Kit for Probes (BioRad, Hercules, CA, US).

### Tissue sampling, histopathology, and *in situ* hybridization

Formaldehyde-fixed bat carcasses were dissected, and organs from both bat species were collected in a standardized manner. To assess the URT, the frontal skull was decalcified for up to 3 weeks and subsequently trimmed in four defined transverse levels, as described [38,39]. These levels permitted evaluation of respiratory epithelium-lined nasal turbinates, predominantly represented in levels I and II, and olfactory epithelium-lined turbinates, predominantly represented in levels III and IV, as well as the nasopharyngeal meatus (S1 Fig). For semiquantitative analysis, turbinate regions were classified according to their predominant epithelium rather than according to individual anatomically defined nasal conchae. Accordingly, the terms “respiratory turbinates” and “olfactory turbinates” are used throughout this study as operational histological categories referring to turbinate mucosa predominantly lined by respiratory or olfactory epithelium, respectively. Because both epithelial types partially overlapped within individual levels, particularly levels II and III, lesion scores and bat H9-specific viral RNA scores were assessed separately for respiratory turbinates, olfactory turbinates, and the nasopharynx. Further specimens from the URT included the palatine tonsil (S2 Fig A), and the larynx.

The LRT was sampled extensively and included the trachea, tracheobronchial lymph nodes (tracheobronchial LNs), right cranial lung lobe, left cranial lung lobe, right caudal lung lobe, and left caudal lung lobe. Samples from the digestive and associated upper alimentary tract included the salivary gland, esophagus, stomach, proximal small intestine, mid small intestine, distal small intestine, cecum, colon, and mesenteric lymph nodes. Additional samples included spleen, liver, and kidney.

To assess potential viral spread to the brain via the olfactory route, brains were sectioned longitudinally to provide an anatomical overview of the olfactory bulb, cortex, striatum, pallidum, thalamus, hippocampus, midbrain, cerebellum, and pons/medulla, as previously described [44].

All tissue samples from all 36 bats were trimmed, paraffin-embedded, sectioned at 2–3 µm on a microtome, and stained with hematoxylin and eosin (H&E) for histopathologic evaluation. Lesion severity was scored on an ordinal scale as follows: 0, no lesion; 1, minimal, affecting <5% of cells or tissue area on the section; 2, mild, affecting 6%–40%; 3, moderate, affecting 41%–80%; and 4, severe, affecting >80%, as previously described [14].

Consecutive tissue sections from all above-mentioned organs of both bat species euthanized at 4 dpi were subjected to in situ hybridization (ISH) using a bat HA-specific vRNA probe to detect viral RNA in tissues. ISH was performed with the RNAscope 2.5 HD Reagent Kit-RED (ACD Bio) in combination with the HybEZ II Hybridization System (ACD Bio) and a 20ZZ probe design. Fast Red served as chromogen, and hematoxylin was used as counterstain. The following probe was used for viral RNA detection: RNAscope Probe - V-InfluenzA-H9N2-segment4-HAC1, targeting IAV (A/bat/Egypt/381OP/2017 [H9N2]) segment 4 hemagglutinin (HA), corresponding to base pairs 35–1275 of GenBank accession no. MH376902.1 (catalog no. 1303251-C1). For both bat species, a negative control probe was included for each individual animal in every run (RNAscope Negative Control Probe, DapB, Bacillus subtilis strain; catalog no. 310043). A positive control probe targeting Rae-PPIB (predicted *Rousettus aegyptiacus* peptidylprolyl isomerase B [PPIB]; catalog no. 573731) was also included in each run and cross-reacted with tissues from *Eidolon helvum*. The distribution of bat H9-specific vRNA was scored semi-quantitatively on an ordinal 0–4 scale as follows: 0, negative; 1, focal or oligofocal; 2, multifocal; 3, coalescing; and 4, diffuse distribution of vRNA-positive cells, as previously described [14]. Histopathologic and ISH evaluation was performed by a board-certified pathologist (JS, Diplomate ACVP).

Chromogenic brightfield images were acquired with a research microscope (Olympus BX53; Olympus Life Sciences, Japan) equipped with an 8.9-megapixel digital camera (DP28-CU; Olympus) and the manufacturer’s software (cellSens Dimension) with 40× and 100×/oil objectives. Whole-slide images were generated using a slide scanner (Axioscan 7; Carl Zeiss Microscopy, Jena, Germany) equipped with a 20× objective (Plan-Apochromat 20×/0.45; Carl Zeiss Microscopy). Photomicrographs were adjusted for white balance, brightness, contrast, and detail sharpening was applied using the unsharp mask filter in Photoshop 2026 (Adobe Inc., San Jose, CA, USA). Supplemental figure 1A was created with ChatGPT.

### Serology

Serological analysis of blood samples from all animals at respective timepoints was performed by an in-house ELISA using the recombinant bat-H9 and bat-N2 proteins as antigens. 100 ng of bat-H9, and 200 ng of bat-N2 per well were coated overnight in 0.1M carbonate buffer pH 9.6 overnight at 4 °C on 96-well microplates (PS, F-Bottom, clear, Microlon, high-binding – Greiner, Frickenhausen, Germany). Next day, plates were washed three times with TBS-T (Sigma-Aldrich, Steinheim, Germany) and blocked with 150 µL 5% milk (Hobbybäcker-Versand, Bellenberg, Germany)) in PBS per well for 1h at 37 °C. Subsequently, plates were washed again three times with TBS-T, and 50 µL of serum samples (1:10 diluted) was added to the appropriate wells, following another incubation period for 1h at 37 °C. Afterwards, cells were washed again three times with TBS-T, following addition of 50 µL multi-species conjugate (SBVMILK; obtained from ID Screen® Schmallenberg virus Milk Indirect ELISA; IDvet, Montpellier,) diluted 1:80 in TBS-T for 1h at room temperature (RT). Plates were washed again three times with TBS-T before initiating the enzymatic color reaction with 100 µL of TMB-substrate for 5-10 minutes. stop solution according to manufacturer’s instructions (IDEXX, Kornwestheim, Germany). Optical density (OD) was measured using 450 nm on a Tecan Infinite® M200 PRO plate reader (Tecan, Männedorf, Switzerland).

Virus neutralizing antibodies in animals that stood out to be seropositive via the bat H9N2-specific ELISA were determined in a live virus neutralization test (VNT_100_) against bat H9N2. Briefly, MDCKII-cells were seeded and grown in 96-well plates for 24 hours. Next day, serum samples were serially diluted in DMEM containing 1 µg per mL TPCK-treated trypsin, following addition and mixing with 100 TCID_50_/mL of live bat H9N2 IAV. Incubation of the virus-serum mix was performed for 2h at 37 °C. Subsequently, the mixture was applied onto the 96-well plates containing the pre-seeded cells. Neutralizing activity was read out 72h after incubation and was considered as neutralizing in the absence of a specific cytopathic effect. The virus neutralizing titer was determined from the last serum dilution in which no CPE was observed.

## Acknowledgments

We thank Angele Breithaupt for performing necropsy of pigs and tissue sampling. We thank Hilke Gräfe of the Institute of Veterinary Pathology for performing in situ hybridization. We gratefully thank Mareen Grawe for the dedicated support in PCR and serological analysis. The authors thank the animal care personnel Frank Klipp, Felix Zimak, Christian Lipinski, Steffen Kiepert, Harald Manthei and Doreen Fiedler for their diligent husbandry and welfare support of the bats in the bat breeding facility, as well as of the pigs and quails, throughout the course of the animal trial. We thank the University Leipzig BioImaging Core Facility (BCF equipment INST 268/230-1; INST 268/293-1; SFB-TR67; EFRE 100192650, 100195814, 100144684) for assistance. We used DeepL (https://www.deepl.com) and GPT-5.6 Thinking (https://chatgpt.com) to improve English-language writing.

---

**Supplement Figure 1.**
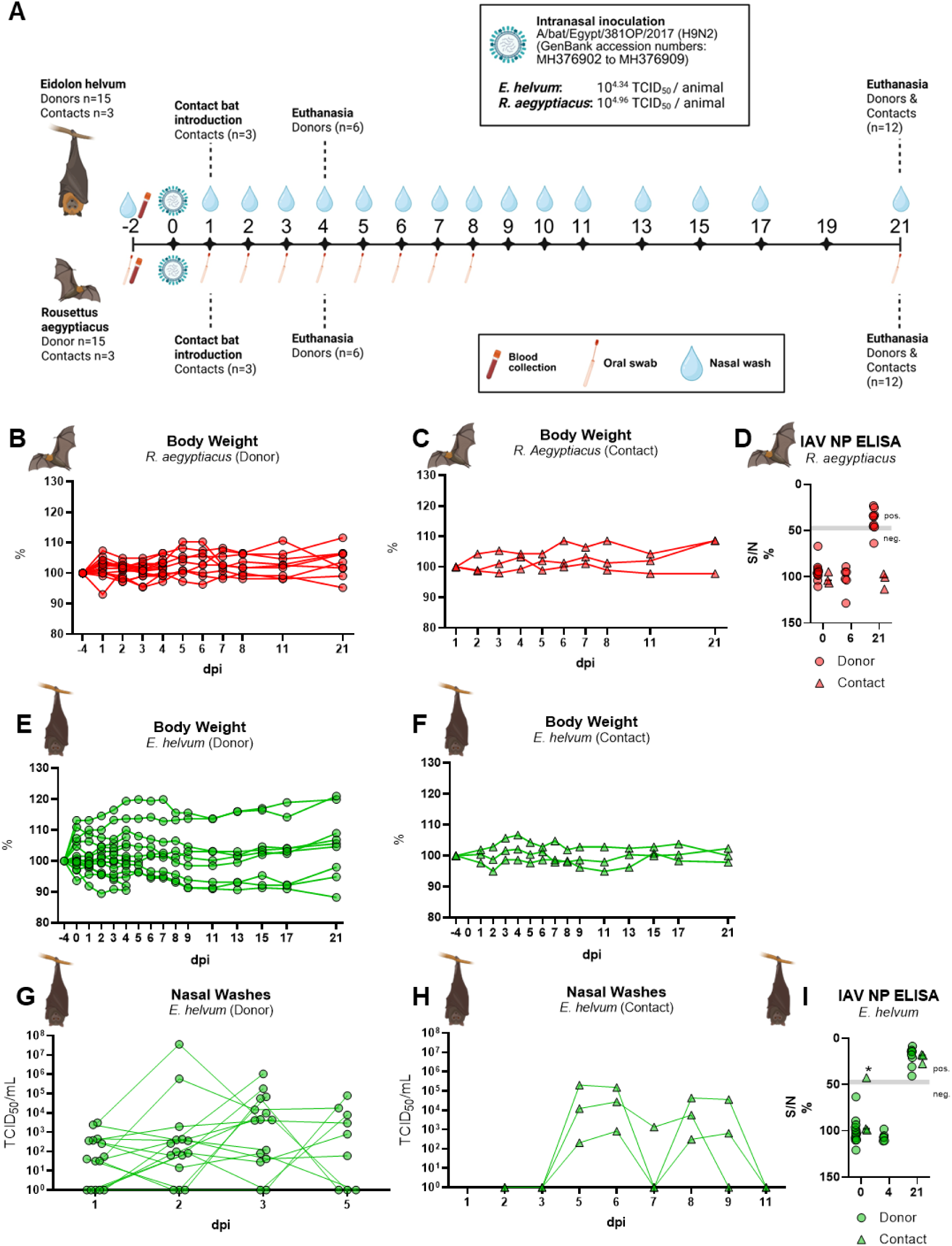
Experimental setup, clinical outcome and IAV-NP ELISA data of bat H9N2 infected *R. aegyptiacus* and *E. helvum.* **(A)** Fifteen bats per species each were experimentally infected with 10^4.34^ or 10^4.9^ TCID_50_ per animal, respectively, of A/bat/Egypt/381OP/2017 (bat H9N2, GenBank accession numbers MH376902 – MH376909). At 1 dpi, 3 contact bats were introduced (donor 15:3 contact setup, one stable per experiment). At 6 dpi (*R. aegyptiacus*) or 4 dpi (*E. helvum*), 6 donor bats were euthanized to study the virus distribution in organ samples. *E. helvum* were sampled daily via nasal washings until 11 dpi, and afterwards every two days until 17 dpi and at the endpoint of the animal trial at 21 dpi. *R. aegyptiacus* were sampled via oral swabs until 8 dpi and at the endpoint of the animal trial at 21 dpi. Performing nasal washing samples in *R. aegyptiacus* was not possible due to their small size. Blood samples were taken two days before and at respective euthanasia timepoints. **(B)** Relative body weight in percent of bat H9N2-infected *R. aegyptiacus* and **(C)** direct contact animals. **(D)** *R. aegyptiacus* serum samples of 0, 6 and 21 dpi were evaluated by IAV NP-specific ELISA. **(E)** Relative body weight in percent of bat H9N2-infected *E. helvum* and **(F)** direct contact animals. **(G)** Infectious virus (TCID50/mL) in 1, 2, 3 and 5 dpi nasal washing samples was determined by endpoint dilution assay of donor animals and **(H)** of contact animals. **(I)** *E. helvum* serum samples of 0, 6 and 21 dpi were evaluated by IAV NP-specific ELISA. An asterisk indicates samples that required predilution because of the very low sample volume.

**Supplement Figure 2.**
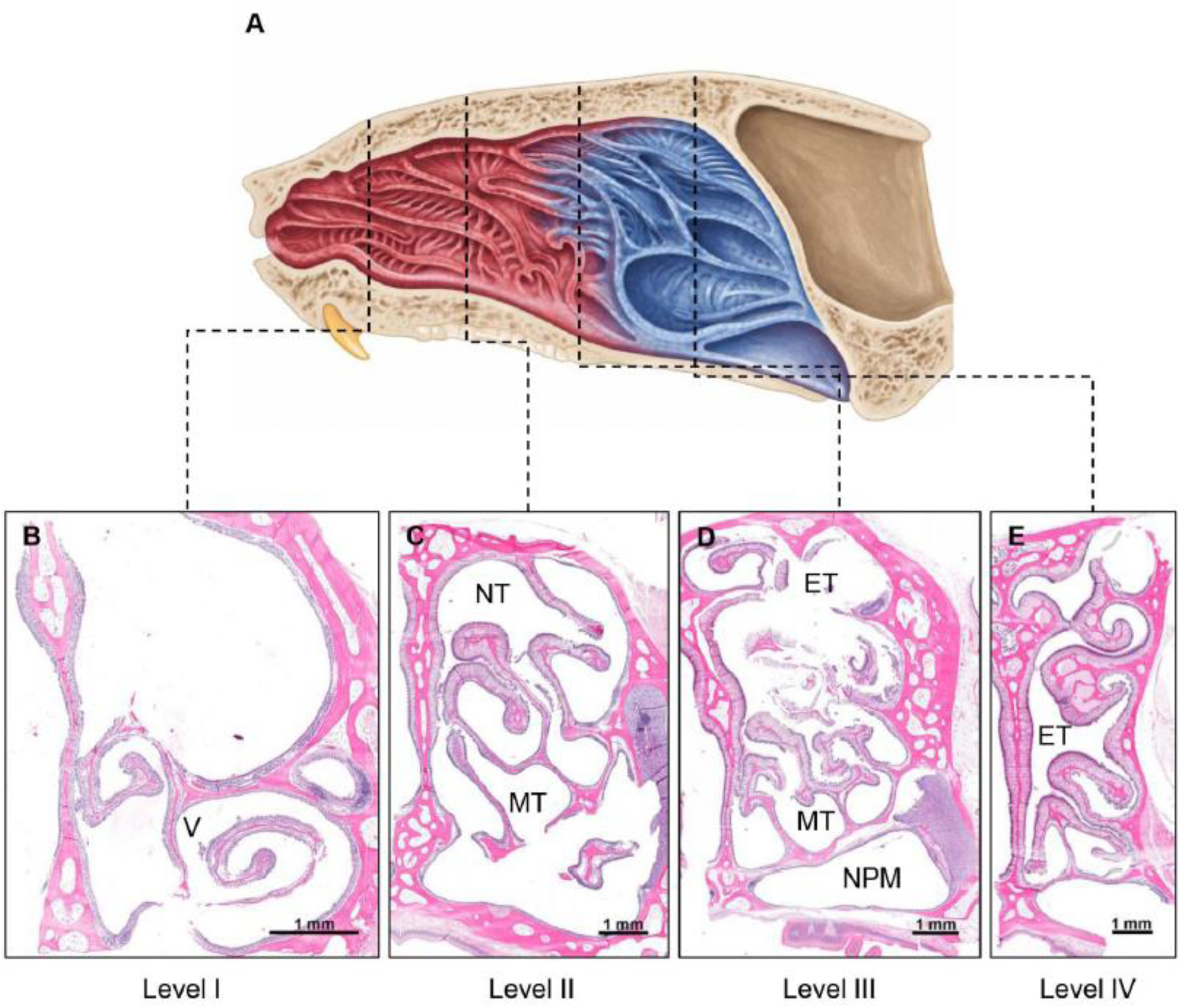
Schematic overview of the bat frontal skull and nasal cavity. **(A)** The rostral nasal cavity is dominated by respiratory turbinates (red), whereas olfactory turbinates (blue) emerge sparsely at mid-levels and become increasingly prominent in the caudal nasal cavity. **(B)** Level I is composed entirely of nasal turbinates lined by respiratory epithelium. **(C)** Level II is still dominated by respiratory epithelium, with only a small extension of olfactory turbinate tissue around the nasoturbinate (NT). **(D)** At Level III, olfactory epithelium predominates, while only minor areas of respiratory epithelium remain at the maxilloturbinate (MT). At this level, the prominent nasopharyngeal meatus, lined exclusively by respiratory epithelium, also becomes visible. **(E)** Level IV shows the ethmoturbinates, which are lined exclusively by olfactory epithelium. V, ventral meatus; MT, maxilloturbinate; NT, nasoturbinate; ET, ethmoid turbinate; NPM, nasopharyngeal meatus.

**Supplement Figure 3.**
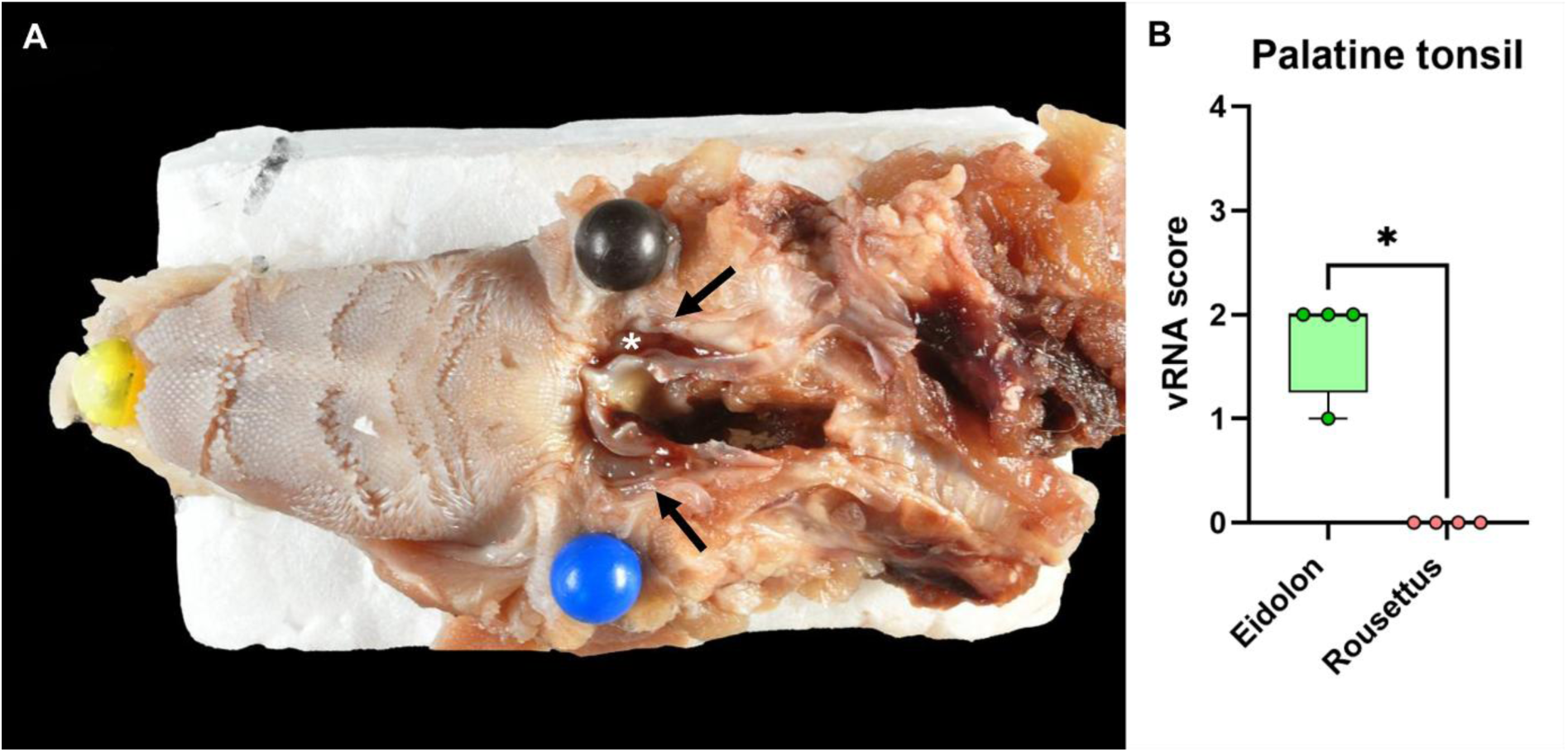
Anatomical location of the palatine tonsils and bat H9-specific vRNA detection by in situ hybridization of *E. helvum* at 4 dpi. **(A)** The paired palatine tonsils (white asterisk) are located deep within a groove adjacent to the epiglottis and are covered by pharyngeal mucosa (arrows). To expose the tonsils for documentation, the overlying tissue was reflected using needles and the preparation was fixed onto cardboard. **(B)** The palatine tonsil shows significantly higher bat H9-specific vRNA scores in *E. helvum* than in *R. aegyptiacus*. Box-and-whisker plots show the median, interquartile range, and minimum-to-maximum range; individual symbols represent single animals (n=4 per group). Statistical comparisons were performed using a two-tailed Mann–Whitney U test. * = p < 0.05.

**Supplement Figure 4.**
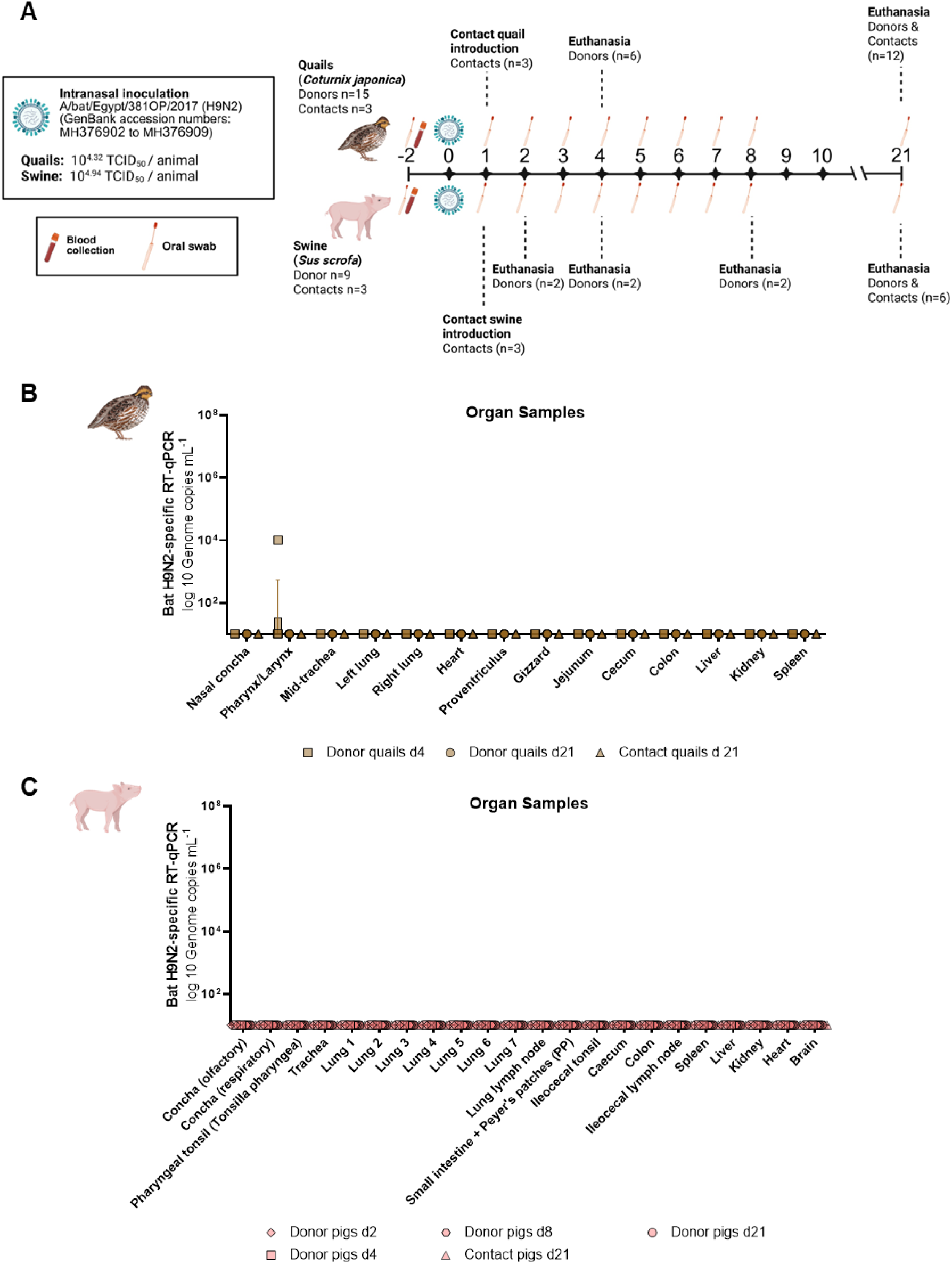
Experimental setup and viral organ load of bat H9N2 infection experiment in quails (*Coturnix japonica*) and pigs (*Sus scrofa dom.*). **(A)** In vivo experimental bat H9N2 infection setups with pigs and quails, carried out independently. 15 quails were experimentally infected with 10^4.94^ TCID_50_ per animal, and 9 pigs were intranasally inoculated with 10^4.32^ TCID_50_ per animal of A/bat/Egypt/381OP/2017 (bat H9N2). 1 dpi, 3 contact quails were introduced to the donor quails (donor 15:3 contact setup in one stable), and 3 contact pigs were introduced to the donor pigs (donor 3:1 contact setup per stable). **(B)** Bat H9N2-specific RT-qPCR analysis of 4 and 21 dpi organ samples of donor and contact quails. **(C)** Bat H9N2-specific RT-qPCR analysis of 2, 4, 8 and 21 dpi organ samples of donor and contact pigs.

